# Horizontal transfer of accessory chromosomes in fungi – a regulated process for exchange of genetic material?

**DOI:** 10.1101/2024.11.21.624634

**Authors:** Michael Habig, Satish Kumar Patneedi, Remco Stam, Henrik Hjarvard De Fine Licht

**Affiliations:** Fungal Evolutionary Genetics, Christian-Albrechts University of Kiel, Germany; Phytopathology and Crop Protection, Christian-Albrechts University of Kiel, Germany; Section for Organismal Biology, Department of Plant and Environmental Sciences, University of Copenhagen, Denmark

## Abstract

Horizontal transfer of entire chromosomes has been reported in several fungal pathogens, often significantly impacting the fitness of the recipient fungus. All documented instances of horizontal chromosome transfers (HCTs) showed a marked propensity for accessory chromosomes, consistently involving the transfer of an accessory chromosome while other chromosomes were seldom, if ever, co-transferred. The mechanisms underlying HCTs, as well as the factors regulating the specificity of HCTs for accessory chromosomes, remain unclear. In this perspective, we provide an overview of the observed propensity in reported cases of horizontal chromosome transfers. We hypothesize the existence of a signal that distinguishes mobile, i.e., horizontally transferred, accessory chromosomes from the rest of the donor genome. Recent findings in *Metarhizium robertsii* and *Magnaporthe oryzae*, suggest that a mobile accessory chromosome may contain putative histones and/or histone modifiers, which could generate such a signal. Based on this, we propose that mobile accessory chromosomes may encode the machinery required for their own horizontal transmission, implying that HCT could be a regulated process. Finally, we present evidence of substantial differences in codon usage bias between core and accessory chromosomes in 14 out of 19 analysed fungal species and strains. Such differences in codon usage bias could indicate past horizontal transfers of these accessory chromosomes. Interestingly, HCT was previously unknown for many of these species, suggesting that the horizontal transfer of accessory chromosomes may be more widespread than previously thought and, therefore, an important factor in fungal genome evolution.

---

Horizontal (lateral) gene transfer (HGT) is characterized as “the movement of genetic information across normal mating barriers between more or less distantly related organisms”(Keeling and Palmer, 2008). In the past decades, the discovery of widespread HGT has completely changed our view of prokaryote evolution, where in some species up to 80% of the genes are believed to have been horizontally transferred (Dagan *et al*., 2008). HGT in eukaryotes, is less common, its importance still being discussed but has been documented in plants (Li *et al*., 2014; Wickell and Li, 2020; Pereira *et al*., 2023), animals (Gladyshev *et al*., 2008; Graham *et al*., 2008; Moran and Jarvik, 2010), and fungi (Slot and Rokas, 2011; Mehrabi *et al*., 2011; Richards *et al*., 2011; Fitzpatrick, 2012). In fungi, HGT can have a crucial role in adaptation and, in some cases, drive significant lifestyle transitions and in some major plant pathogens affect virulence (Friesen *et al*., 2006; Keeling and Palmer, 2008; Fitzpatrick, 2012; McDonald *et al*., 2018; Zhang *et al*., 2019; Etten and Bhattacharya, 2020; Sahu *et al*., 2023; Ciach *et al*., 2024). Exactly how such HGTs occur in fungi is poorly understood but one avenue appears to be through horizontal chromosome transfer (HCT). HCT involves transmission of entire chromosomes from a donor to a recipient, which represents a massive HGT event potentially involving hundreds of genes located on the transmitted chromosome. In fungi, such HCTs have been reported for some accessory chromosomes, which are chromosomes that show presence/absence polymorphism. This apparent specificity of HCT for accessory chromosomes might be indicative of a regulated mechanism, possibly by the transferred accessory chromosome itself, which would be conceptually similar to conjugative bacterial plasmids, which encode the machinery for their own transfer and represent one of the pivotal avenues for HGT in prokaryotes.

## Accessory Chromosomes are found in several, mostly pathogenic fungi, often affecting their pathogenicity and/or host-range

Accessory chromosomes are nonessential, linear chromosomes found in some but not all members of a population. Also known as B chromosomes, supernumerary chromosomes, dispensable chromosomes, lineage-specific chromosomes, or mini-chromosomes, they have been reported in over 3,000 eukaryotic species, including plants, animals, and fungi (Houben, 2017; D’Ambrosio *et al*., 2017; Ferree *et al*., 2024). Many fungal species, particularly pathogenic fungi, possess distinct accessory chromosomes that can constitute a significant portion of their genomes. Overall, accessory chromosomes have been reported in more than 25 fungal species, including major plant pathogens like *Magnaporthe oryzae, Fusarium oxysporum, Alternaria alternata, F. solani* (synonym *F. vanettinii*), and *Zymoseptoria tritici* (Orbach *et al*., 1996; Han *et al*., 2001; Coleman *et al*., 2009; Ma *et al*., 2010; Goodwin *et al*., 2011; Langner *et al*., 2021). Accessory chromosomes in pathogenic fungi often carry virulence factors directly linked to pathogenicity. They can affect the host range of the fungus, for instance, the *SIX* effectors located on the accessory chromosome 14 of the plant pathogen *F. oxysporum* f.sp. *lycopersici* are required for its pathogenicity on tomato (Ma *et al*., 2010; Hu *et al*., 2012; Zhang *et al*., 2020). Another example are the accessory chromosomes of *A. alternata* which encode host specific toxins (Akamatsu *et al*., 1999; Johnson *et al*., 2001). Accessory chromosomes are defined by their presence/absence polymorphism, which can be affected by their segregation during mitotic and meiotic cell-divisions (Komluski *et al*., 2022). Readers are directed to comprehensive reviews on accessory chromosomes in fungi, which discuss their prevalence, potential function, their presence-absence polymorphism and unique characteristics (Mehrabi *et al*., 2011, 2017; Bertazzoni *et al*., 2018; Habig and Stukenbrock, 2020). We here focus on their potential for horizontal transmission – which is distinct from their vertical transmission during canonical mitosis and meiosis.

## Accessory chromosomes are horizontally transferred in some fungi

Fungal accessory chromosomes can be horizontally transferred, representing a significant horizontal gene transfer (HGT) event, often encompassing hundreds of genes and, in addition, non-genic elements. Horizontal transfer of entire accessory chromosomes has been experimentally documented in a number of fungi, including *F. oxysporum* (Ma *et al*., 2010; van Dam *et al*., 2017; Li *et al*., 2020), *Alternaria* spp. (Akagi *et al*., 2009), *Metarhizium robertsii* (Habig *et al*., 2024), and *Colletotrichum gloeosporioides* (He *et al*., 1998). In addition to experimental evidence, phylogenetic analyses provide support for horizontal transfer events of accessory chromosomes in further *F. oxysporum* species and strains (Ma *et al*., 2010; van Dam *et al*., 2017), *Alternaria* spp. (Wang *et al*., 2019), *M. oryzae* (Barragan *et al*., 2024) and between *Metarhizium* spp (Habig *et al*., 2024). It is important to note that HCT appears to not require meiosis and is limited to specific chromosomes (see next paragraph). This makes it distinct from (sexual) hybridzation, which is already recognized as an important factor in fungal evolution (Steensels *et al*., 2021).

## Reported instances of horizontal chromosome transfer (HCT) in fungi are not random but show a clear pattern of preference and specificity for accessory chromosomes

What could be a mechanism for HCT? Entire eukaryotic chromosomes are likely not stable outside cells and therefore cytoplasmic continuity - either between hyphae or germ tubes (Roca *et al*., 2004; Fleissner *et al*., 2009; Ishikawa *et al*., 2010) - is believed to be a prerequisite for horizontal chromosome transfer (Mehrabi *et al*., 2011). Following such cellular plasmogamy, heterokaryons can progress to parasexuality, a mechanism observed in some fungi that allows for the recombination of genetic material independent of meiosis (Fig. 1) (Nieuwenhuis and James, 2016; Yadav *et al*., 2023). In this process, the ploidy of the fused nucleus (now diploid) is reduced back to haploidy through presumably random chromosome loss and possibly involving mitotic recombination. Consequently, parasexuality is considered to lead to a random reassortment of genetic material from the two fused cells. In stark contrast, none of the reported instances of fungal HCT demonstrate random reassortment of genetic material; instead, there is a marked specificity, i.e. preference for particular accessory chromosomes, even when other accessory chromosomes are also present. Furthermore, HCT very rarely involves core chromosomes. Here, we provide a brief overview of the examples of HCT in fungi, emphasizing the aspect of this specificity.

**Fig. 1:**
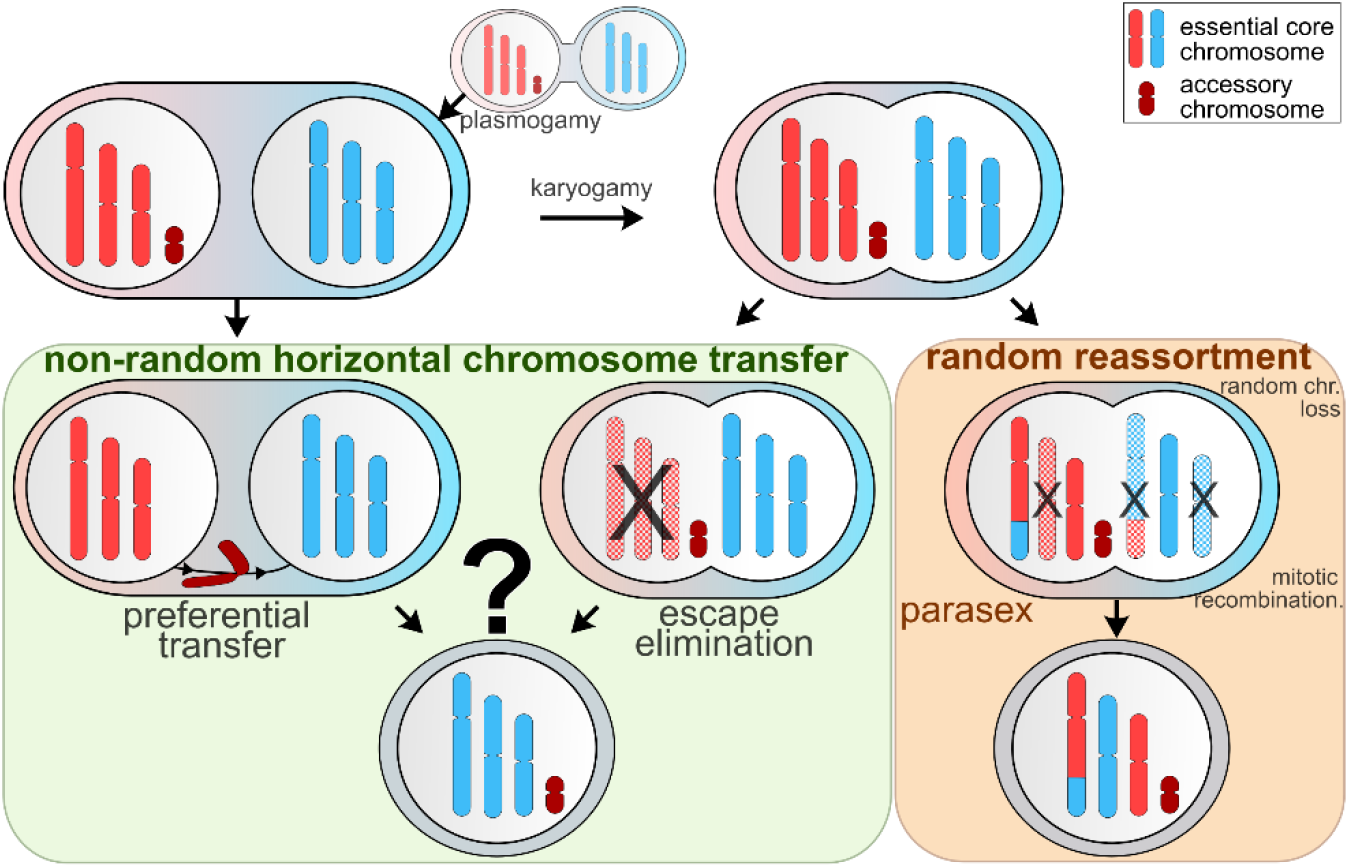
Proposed mechanism for non-random horizontal chromosome transfer, contrasted with the expected random chromosome reassortment that occur during parasexuality. First, two fungal cells or hyphae must anastomose so divergent nuclei can interact (if two haploid nuclei are involved this can lead to plasmogamy). Second, this may or may not lead to nuclear fusion (karyogamy). Left, green box: The non-random horizontal transfer of accessory chromosomes could involve preferential transfer between nuclei (or from a degraded nucleus to an intact one). Alternatively, if karyogamy occurs, there may be selective retention of the accessory chromosome while the remaining donor chromosomes are eliminated (as suggested by (He *et al*., 1998; Vlaardingerbroek *et al*., 2016)). Both mechanisms would require a signal that differentiates the accessory chromosome from the rest of the genome. Right, brown box: In contrast, during fungal parasexuality, following karyogamy, the diploid nucleus can revert to haploidy through subsequent, presumably random chromosome loss and in addition mitotic recombination may occur, resulting in random chromosome reassortment.

In *A. alternata*, induced protoplast fusion resulted in horizontal chromosome transfer (HCT) of an accessory chromosome, referred to as a conditionally dispensable chromosome in this fungus. Although complete genome sequencing data were not generated, there was no indication that any other genetic material was transferred from the donor to the recipient (Akagi *et al*., 2009). Similarly, in *C. gloeosporioides*, HCT of an 2 Mb accessory chromosome was observed between two vegetatively incompatible biotypes, involving only the accessory chromosome while no other genetic material was transferred from the donor to the recipient. Additionally, attempts to detect the transfer of core chromosomes using the same system of different fungicide resistance markers in donor and recipient strains were unsuccessful for these biotypes further indicating that the ability of horizontal transfer appears to be restricted to the accessory chromosome (He *et al*., 1998).

Members of the *Fusarium oxysporum* species complex (FOSC) have been most intensively studied for HCT. In this group, accessory chromosomes—referred to as lineage-specific chromosomes—are mobile and capable of horizontal transfer. Accessory chromosome 14, often called the pathogenicity chromosome, along with parts of accessory chromosomes 3 and 6, has been shown to be horizontally transferred from *F. oxysporum* f.sp. *lycopersici*, which is pathogenic on tomato, to a non-pathogenic strain, thereby rendering the recipient strain pathogenic on tomato as well. Importantly, no core chromosomes were transferred in this process (Ma *et al*., 2010). In a later study, again only accessory chromosome 14 was horizontally transferred, while other accessory chromosomes, albeit also being targeted and analysed similar to chromosome 14, were not transferred. In this study, parts of core chromosomes were also transmitted, but these transfers were always accompanied by the transfer of accessory chromosome 14, even though in these cases it did not carry the fungicide resistance marker that was used to detect HCT (Vlaardingerbroek *et al*., 2016). Similarly, in *F. oxysporum* f.sp. *radicis-cucumerinum*, an accessory chromosome was repeatedly transferred horizontally, while no other donor genetic material was transferred (van Dam *et al*., 2017). In summary, HCT has been frequently reported in FOSC, specifically involving one accessory chromosome, although transfers of core chromosomal material can occur in rare cases.

Very recently, horizontal chromosome transfer (HCT) of accessory chromosomes has been reported in the entomopathogenic *Metarhizium robertsii* and in *Magnaporthe* (syn. *Pyricularia*) *oryzae*. In *M robertsii*, an accessory chromosome (termed chrA) was frequently horizontally transmitted from a donor to a recipient strain, while the transfer of any other genetic material could be ruled out with high certainty based on long- and short-read sequencing data (Habig *et al*., 2024). This transfer occurred during co-infections of a host insect by different strains of *M. robertsii*, and apparently affected the fitness of the recipient fungus. Notably, additional phylogenetic analysis showed that chrA had also been horizontally transmitted between distinct *Metarhizium* species. In addition, in *M. oryzae*, a plant pathogen responsible for crop pandemics through asexual reproduction of clonal lineages, accessory chromosomes (referred to as mini-chromosomes in this fungus) contain virulence-related genes and have been horizontally transferred. In particular, the accessory chromosome mChrA—a 1.2 Mb mini-chromosome containing 191 genes—has been frequently horizontally transferred between clonal lineages of *M. oryzae* in the recent past. Again, notably, there was no indication of core genome exchange (Langner *et al*., 2021; Barragan *et al*., 2024).

In summary, all reported horizontal chromosome transfers showed a high degree of propensity and specificity towards involving a specific accessory chromosome and hence cannot be solely explained by the assumingly random processes of parasexuality (Fig. 1).

## The non-random process of horizontal chromosome transfer (HCT) requires a distinguishing signal that differentiates mobile chromosomes from the rest of the genome

The non-random nature of HCT has already been previously acknowledged, and two primary mechanisms causing this specificity have been proposed (He *et al*., 1998; Vlaardingerbroek *et al*., 2016). The first suggests that the mobile chromosome is preferentially transferred during the heterokaryon stage from one (potentially degrading) donor nucleus to the recipient nucleus (Fig. 1). The second mechanism posits that following karyogamy, the mobile chromosome avoids a targeted genome elimination process (Fig. 1). A related phenomenon to the former has been observed in the budding yeast *Saccharomyces cerevisiae*, where a mutant with defective nuclear fusion temporarily create heterokaryons, which facilitate chromosome transfer between nuclei before one of them is lost – a process termed chromoduction (Conde and Fink, 1976; Dutcher, 1981; Ji *et al*., 1993; Zhao *et al*., 2022). Similarly, a process akin to the second mechanism—where all but the mobile chromosomes are degraded—occurs during programmed DNA elimination observed in various plants and animals (Dedukh and Krasikova, 2022). Both mechanisms require a signal that distinguishes mobile chromosomes from the rest of the genome, either for preferential transfer or to avoid elimination. We speculate such a signal could be epigenetic and we base our speculation on previous examples of histone modification-regulated genome elimination, such as in the jewel wasp *Nasonia vitripennis* (Aldrich *et al*., 2017; Lee *et al*., 2023), and the distinct epigenetic modifications that differentiate fungal accessory chromosomes from core chromosomes (Schotanus *et al*., 2015; Freitag, 2017; Habig *et al*., 2021).

## Could mobile accessory chromosomes encode the functions for their own horizontal transfer?

Functional annotation of genes on the mobile accessory chromosomes chrA and chrB in *M. robertsii* revealed an enrichment of genes that potentially influence chromatin conformation (Habig *et al*., 2024). Notably, chrA harbors two putative histone H2B genes, two putative histone lysine N-methyltransferases (H3 lysine-4 specific) (Freitag, 2017), and a putative histone acetyltransferase similar to GCN5 (predominantly H3 lysine 14 specific) (Li and Shogren-Knaak, 2009). Importantly, these genes are not paralogs of their canonical counterparts on core chromosomes but true xenologs, suggesting they may exhibit different specificities. Similarly, the mobile accessory chromosome mChrA in *M. oryzae* is enriched (among others) for genes involved in the GO-terms for the molecular function of chromatin binding and for genes encoding products located in the nucleosome (Barragan *et al*., 2024). Moreover, mChrA of *M. oryzae* also contains genes encoding two putative histones (g11257 and g11315) (Barragan *et al*., 2024). The presence of these potentially chromatin-affecting genes on the mobile accessory chromosome chrA in *M. robertsii* as well as the mobile accessory chromosome mchrA in *M. oryzae* raises the intriguing possibility that these mobile chromosomes encode functions necessary for their own preferential transfer, conceptually similar to conjugative plasmids in bacteria, which also encode the machinery for their own horizontal transfer. Given that HCT of fungal accessory chromosomes in other species also display some degree of specificity, these mobile chromosomes could potentially also be involved in their own transmission as well. However, genes located on fungal accessory chromosomes are poorly annotated functionally, with few characterized beyond putative effectors. For instance, in *A. alternata*, more than 25 genes identified on an accessory chromosome of multiple isolates lack any functional annotation, and most appear to be unique to the *Alternaria* genus. (Harimoto *et al*., 2007; Gai *et al*., 2021). Functionally annotating these and other genes on accessory chromosomes could be highly insightful, potentially uncovering functions that, for example, influence chromatin conformation. These would be intriguing targets for testing their impact on the horizontal transfer rates of these chromosomes.

## How widespread is horizontal chromosome transfer of accessory chromosomes?

Despite the presence of accessory chromosomes in more than 25 fungal species (Bertazzoni *et al*., 2018; Komluski *et al*., 2022), horizontal transfer has only been reported in five species. Since HCT appears to specifically involve accessory chromosomes, those accessory chromosomes, for which HCT have not been reported yet, might still have experienced HCTs in the past and the consequences of these transfers might still be detectable. One measurable indication of past horizontal transfers is variation in the relative synonymous codon usage (RSCU) between core and accessory chromosomes. RSCU, introduced by Sharp and Li (Sharp and Li, 1986), measures codon usage bias, which can be influenced by mutational processes such as GC content and nucleotide sequence context, as seen in mechanisms like Repeat-Induced Point (RIP) mutation (Galagan and Selker, 2004; Hershberg and Petrov, 2008; Plotkin and Kudla, 2011). Crucially, codon usage bias is also shaped by selective processes, where certain preferred codons are translated more accurately or efficiently (Hershberg and Petrov, 2008). This form of translational selection is thought to correlate with relative tRNA abundance (Plotkin and Kudla, 2011). Thus, both mutational and selective pressures may vary between species and genomes, potentially leading to differences in codon usage bias. Since horizontal chromosome transfer (HCT) involves the movement of an accessory chromosome from one genomic background—where it was shaped by species-specific mutational and selective forces—into a new genomic background where these forces may differ, we hypothesize that HCT might be reflected in differences in the codon usage bias between core and accessory chromosomes (Fig. 2). Indeed differences in codon usage bias between core and accessory chromosomes, had previously been reported for several fungal species, e.g. *F. oxysporum, F. vanettinii* MPVI, *A. arborescens, C. graminicola* and *M. robertsii* (Coleman *et al*., 2009; Ma *et al*., 2010; Goodwin *et al*., 2011; Hu *et al*., 2012; Becerra *et al*., 2023; Habig *et al*., 2024).

**Fig. 2:**
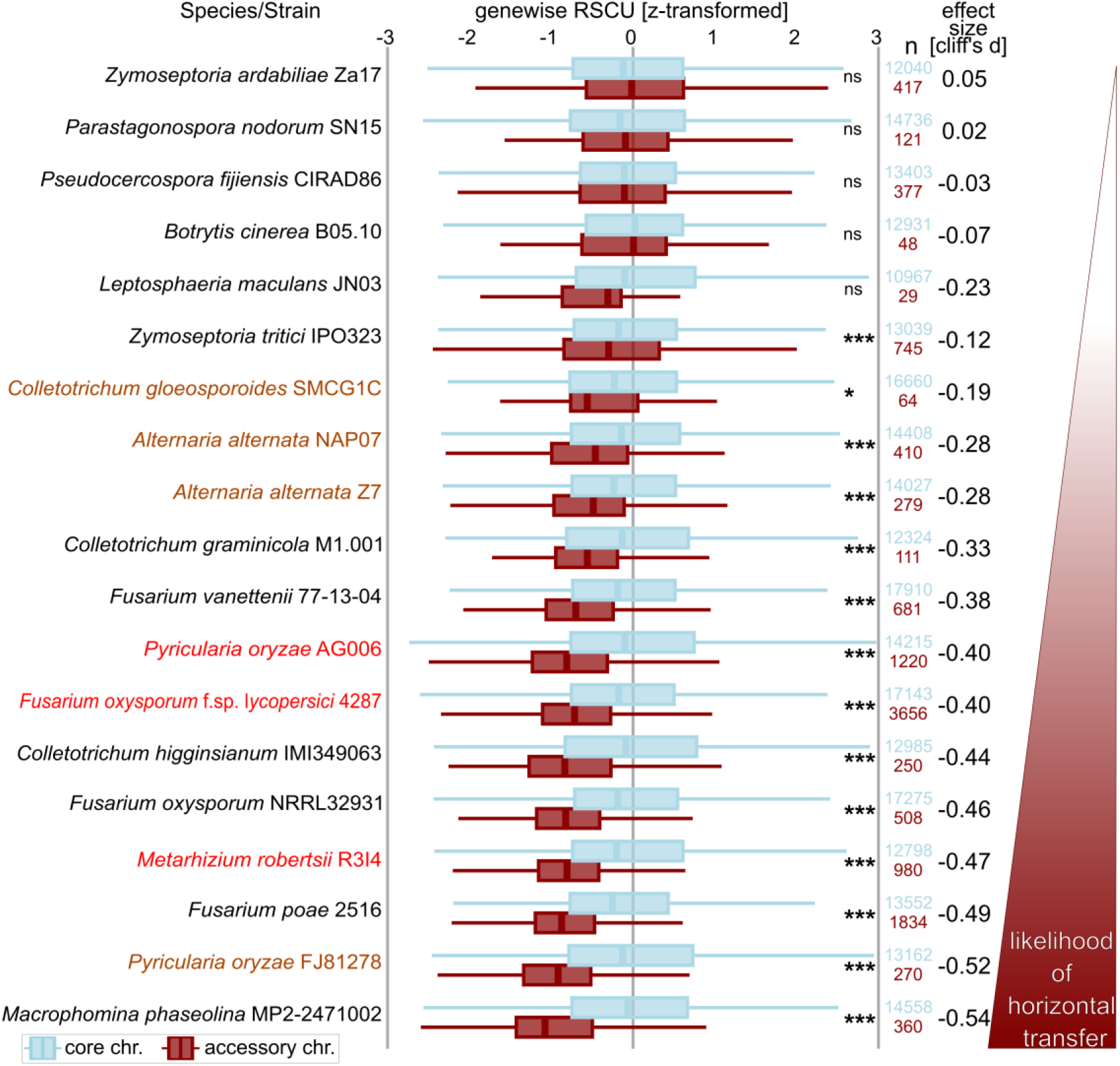
Differences in codon usage bias between core and accessory chromosomes may indicate horizontal transfer. Codon usage bias, assessed through gene-wise RSCU (Steenwyk *et al*., 2022) comparisons between core (blue) and accessory chromosomes (dark red), z-transformed for easier visual comparison (see also Fig S1 for untransformed data). Fungal strains with documented horizontal transfer are highlighted in red, while those with evidence of transfer in different strains of the same species are marked in orange. Species are organized according to increasing effect sizes, represented by the absolute value of Cliff’s d, for those exhibiting significant differences. Statistical significance was determined using the Wilcoxon rank sum test (with Hochberg adjustment for multiple testing). Not significant (ns), * = p < 0.05, ** = p < 0.005, *** = p < 0.0005. n represents the number of cds that were included for core (blue) and accessory chromosomes (red).

However, it is important to note that differences in codon usage bias may also arise from mutational or selective processes that affect accessory chromosomes differently than core chromosomes independently of horizontal transfer. For instance, accessory chromosomes may exhibit higher mutation rates due to distinct histone modifications (Habig *et al*., 2021), or in sexually reproducing organisms, these chromosomes may be subjected to less efficient selection due to their smaller effective population sizes. Additionally, for individual genes, codon usage bias is not a sensitive indicator of past horizontal transfer (Koski *et al*., 2001). Nevertheless, because HCT typically involves the transfer of a large number of genes and non-genic elements, we believe that analysing codon usage bias across all transcripts from core and accessory chromosomes could reveal significant differences that warrant further investigation. It is also important to note that the absence of differences in codon usage bias does not necessarily rule out past horizontal transfers, as HCT could occur between species with similar genome-wide codon usage bias. Therefore, while a pronounced difference in codon usage bias may be a strong indicator of past horizontal transfer, the lack of such differences should not be interpreted as definitive evidence against it. To explore this hypothesis, we analysed codon usage differences between core and accessory chromosomes in fungal species where HCT has been experimentally or phylogenetically demonstrated (Fig. 2). Additionally, we included fungal species with available genome sequencing data that distinguish between core and accessory chromosomes, to explore how widespread signals of potential HCT are among fungal accessory chromosomes. To be able to compare the potential differences between species and strains we annotated the genes *ab initio* for all (see Text S1 for Methods and Materials).

## Horizontally transferred accessory chromosomes exhibit distinct codon usage biases, which is a characteristic shared by many other accessory chromosomes and suggests more widespread horizontal chromosome transfers

By comparing codon usage bias between core and accessory chromosomes across 19 strains from 16 distinct fungal species (see Text S1 for details), we found that five species—*Parastagonospora nodorum, Botrytis cinerea, Pseudocercospora fijiensis*, and *Leptosphaeria maculans*—showed no significant difference in codon usage between core and accessory chromosomes (Fig. 2). Notably, these species are known or assumed to undergo frequent sexual reproduction (Rouxel and Balesdent, 2005; Williamson *et al*., 2007; Churchill, 2010; Stukenbrock *et al*., 2012; Gao *et al*., 2016), and hence the accessory chromosomes might be primarily vertically inherited. Conversely, we found significant and highly pronounced differences (measured by the effect size using Cliff’s delta, a robust non-parametric estimate of effect sizes that does not assume normally distributed data) between core chromosomes, as and accessory chromosomes, for which horizontal transfer had been experimentally or phylogenetically demonstrated. This included *F. oxysporum* f.sp. *lycopersici* strain 4287 (Ma *et al*., 2010), *M. oryzae* strain AG006 (Barragan *et al*., 2024), and *M. robertsii* strain R3I4 (Habig *et al*., 2024) (see Fig. 2 in red). Additionally, significant differences were also detected in strains of species, for which horizontal transfer was experimentally shown in only a different strain than that with sequencing data, including *C. gloeosporioides* (He *et al*., 1998) and *A. alternata* (Akagi *et al*., 2009) (see Fig. 2, orange). Hence, significant and pronounced differences in the RSCU were detectable for all accessory chromosomes for which horizontal transmission was previously shown. For the remaining accessory chromosomes, for which it is unknown whether horizontal transfer occurs, we see a variation from relatively small but significant differences in *Z. tritici* IPO323(Cliff’s d = -0.11), to large significant difference in *Macrophomina phaesolina* strain MP2-2471002 (Cliff’s d= -0.57). In particular the large, significant difference in codon usage bias for the accessory chromosome(s) of *M. phaseolina*, a broad host-range asexual plant pathogen for which parasexuality is suspected (Jones *et al*., 1998; Marquez *et al*., 2021), as well as *F. poae* and *C. higgensianum* is striking and should warrant further investigation into HCT. In general, the fact that the majority of accessory chromosomes show a large significant difference in codon usage could be indicative of many more horizontal chromosome transfers of accessory chromosomes than are currently known.

## Final Remarks

Horizontal chromosome transfer (HCT) has been experimentally and phylogenetically demonstrated in a handful fungal species with accessory chromosomes, often significantly impacting the fitness, pathogenicity, or life-history traits of the recipient fungus. The fact that such HCTs consistently show specificity for accessory chromosomes suggests that it may be a regulated process, potentially enabling the extensive horizontal transfer of both genic and non-genic elements and thereby influencing fungal genome evolution. A simple comparison of relative synonomyous codon usage (RCSU) bias between accessory and core chromosomes suggests that HCT might be much more prevalent among fungal accessory chromosomes than previously thought, highlighting the potential significance of this process in fungal evolution. Mobile accessory chromosomes, with distinct transmission systems compared to core chromosomes, may exhibit separate evolutionary dynamics and potentially act as selfish genetic elements. Unravelling the signals and genetic mechanisms underlying HCT will require a functional dissection of genes located on accessory chromosomes, especially genes that might influence chromatin conformation or chromosome segregation. This focus could be particularly valuable for understanding the unique and impactful phenomenon of horizontal transfer of entire chromosomes.

